# ScRNA-seq Identified the Metabolic Reprogramming of Human Colonic Immune Cells in Different Locations and Disease States

**DOI:** 10.1101/2022.02.14.480452

**Authors:** Qiuchen Zhao, Tong Zhang, Hao Yang

## Abstract

Different regions and states of the human colon are likely to have a distinct influence on immune cell functions. Here we studied the immunometabolic mechanisms for the spatial immune specialization and the dysregulated immune response during ulcerative colitis at single-cell resolution. We revealed that the macrophages and CD8^+^ T cells in the lamina propria (LP) of the human colon possessed an effector phenotype and were more activated, while their lipid metabolism was suppressed compared with those in the epithelial (Epi). Also, IgA^+^ plasma cells accumulated in lamina propria of the sigmoid colon were identified to be more metabolically activated versus those in the cecum and transverse colon, and the improved metabolic activity was correlated with the expression of CD27. In addition to the immunometabolic reprogramming caused by spatial localization colon, we found dysregulated cellular metabolism was related to the impaired immune functions of macrophages, dendritic cells, regulatory T cells, and ILCs in patients with ulcerative colitis. The cluster of OSM^+^ inflammatory monocytes was also identified to play its role in resistance to anti-TNF treatment, and we explored targeted metabolic reactions that could reprogram them to a normal state. Altogether, this study revealed a landscape of metabolic reprogramming of human colonic immune cells in different locations and disease states, and offered new insights into treating ulcerative colitis by immunometabolic modulation.

## INTRODUCTION

The colon, as a barrier tissue, is crucial in immunological research due to its special immune environment (Zhang and Liu, 2016). There are evident regional differences when comparing the epithelial and lamina propria or comparing with different locations of the colon in humans (James et al., 2020a), which are usually reflected in terms of the composition of the immune system, for example, the percentages of immune cells differ from MLNs (mesenteric lymph nodes) to sigmoid (Berthold et al., 2021). And compared to epithelial, lamina propria holds more B cells, macrophages, dendritic cell (Hirata et al., 1986; Panduro et al., 2016; Weigmann et al., 2007), displaying an effector memory phenotype in T cells. (Berthold et al., 2021). The majority of human intraepithelial lymphocytes are T cells, although higher proportions of non-T cells are found within the human colonic epithelium (Selby et al., 1981). Besides, there was a marked increase in the activation and memory status of T and B cells in MLNs compared with the other colon sections (James et al., 2020a), while the colonic CD4^+^ T cells had a more effector phenotype (Kumar et al., 2017), suggesting that these cell types are molded by their environment. In addition to the differences caused by the region of the colon, changes of immune system during colonic disease states are also quite drastic. Some studies focus on the alterations of composition of the immune cells during ulcerative colitis (Smillie et al., 2019c), for example, there are increases in the proportions of mast cells, CD8^+^IL-17^+^ T cells, and Treg cells (Holmen et al., 2006; King et al., 1992; Tom et al., 2016).

Although major immune system conditions have been studied in the mouse intestine, cell type-specific markers and functional assignments are largely unavailable for the human intestine (Wang et al., 2020). In recent years, Single-cell RNA sequencing (scRNA-seq) technology has been greatly developed, which has been widely used in intestinal studies and is helping to enhance our understanding of the human disease effect by comprehensively mapping the cell types and states within the intestines (James et al., 2020c; Mitsialis et al., 2020; Sato et al., 2011; Smillie et al., 2019b). Recent studies in the field of immunometabolism at the single-cell level have demonstrated a close connection between metabolic programs and the specific immune functions they support during both health and disease (Makowski et al., 2020). However, the research on intestinal immunometabolism at single cell resolution is limited nowadays, so it is vital to study immunometabolism by using novel computational algorithms based on single-cell technologies. Currently, the pathway enrichment pipelines can be adjusted for scRNA-seq data analysis and utilized to characterize the metabolic diversity among immune subsets (Xiao et al., 2019), and some systems-based approaches, including Single-cell Flux Balance Analysis (scFBA) (Damiani et al., 2019; Wagner et al., 2021) and FBA-independent approaches (Alghamdi et al., 2021a), were also translated to scRNA-seq data to investigate metabolic differences in immune cells during disease states.

In this article, our study aimed to explore the metabolic reprogramming of immune cells in different regions and states of the human colon at single-cell resolution. To this end, we analyzed colon data in different locations and with disease models of humans trying to investigate the alterations of immunometabolism in the human colon (James et al., 2020a; Smillie et al., 2019c). And the colonic immune systems corresponding to different and the intestine under the disease states have been studied with novel algorithms for single-cell metabolism analysis, which would be helpful for a better understanding of human intestine disorders, such as inflammatory bowel disease and intestinal cancer (Donaldson et al., 2016; James et al., 2020a; Smillie et al., 2019c).

## RESULTS

### Distinct immune response in colonic epithelium versus lamina propria

Based on the questions posed in the introduction, we obtained scRNA-seq data of immune cells (about 260,000 cells) from different locations and disease states of the human colon (James et al., 2020a; Smillie et al., 2019c) and performed single-cell metabolism analysis **(Figure 1A)**. Firstly, we sought to explore the immune heterogeneity between epithelial and lamina propria. The relative percentage of each immune subsets in Epi and LP shows CD8^+^ T cells occupy the largest immune group in Epi, while the plasma and CD4^+^ T cells are the main cell types in the LP **(Figure 1B)**. We then determined the significantly different gene expression between two groups. Surprisingly, nearly all the immune subsets in Epi expressed FABP1 (fatty acid binding protein 1 (McArthur et al., 1999)), while those in LPs did not **(Figure 1C)**. We also found that the macrophages in LP had a relatively suppressed ability of fatty acid degradation and were more activated compared with those in Epi using Gene set enrichment analysis (GSEA) and signature score evaluation **(Figure 1D-E, Figure S1A-B)**, which was consistent with previous finding that FABP1 was positively related to the transport of fatty acid and negatively correlated with the inflammatory response in macrophages (Jin et al., 2021). The genes mattering with immune activation are expressed highly in macrophages of LP, such as IL1B, IL8, CCL3, and NFKBIA (Abplanalp et al., 2021; Chua et al., 2020). In addition to the metabolic gene expression level, we used a mathematical approach termed scFEA (single-cell Flux Estimation Analysis) (Alghamdi et al., 2021a) to analyze the flow of metabolites through a metabolic network. Leveraging the multi-scale stoichiometric models, we found the fatty acid to acetyl-CoA flux was significantly increased in Epi macrophages **(Figure 1G)**. Then we examined the CD8^+^ T cells in Epi and LP similarly, the results of GSEA and scFEA suggested that the CD8^+^ T cell in LP might be more activated, while possessed lower fatty acid metabolism **(Figure 1H-I, 1K)**, which was consistent with results of fatty acid oxidation and CD8^+^ T activation scores **(Figure S1C-D)**. Some key genes related to the CD8^+^ T cell activation were significantly upregulated in the CD8^+^ T cells from LP, such as CCL4, IFNG, NKFBIA, while CD160 which had a negative influence on T cell priming (Liu et al., 2020) was downregulated **(Figure 1J)**. In a nutshell, distinct immune response was explored in colonic epithelium versus lamina propria, and colonic immune cells in lamina propria were more activated with reduced activity of fatty acid metabolism.

**Figure 1.**
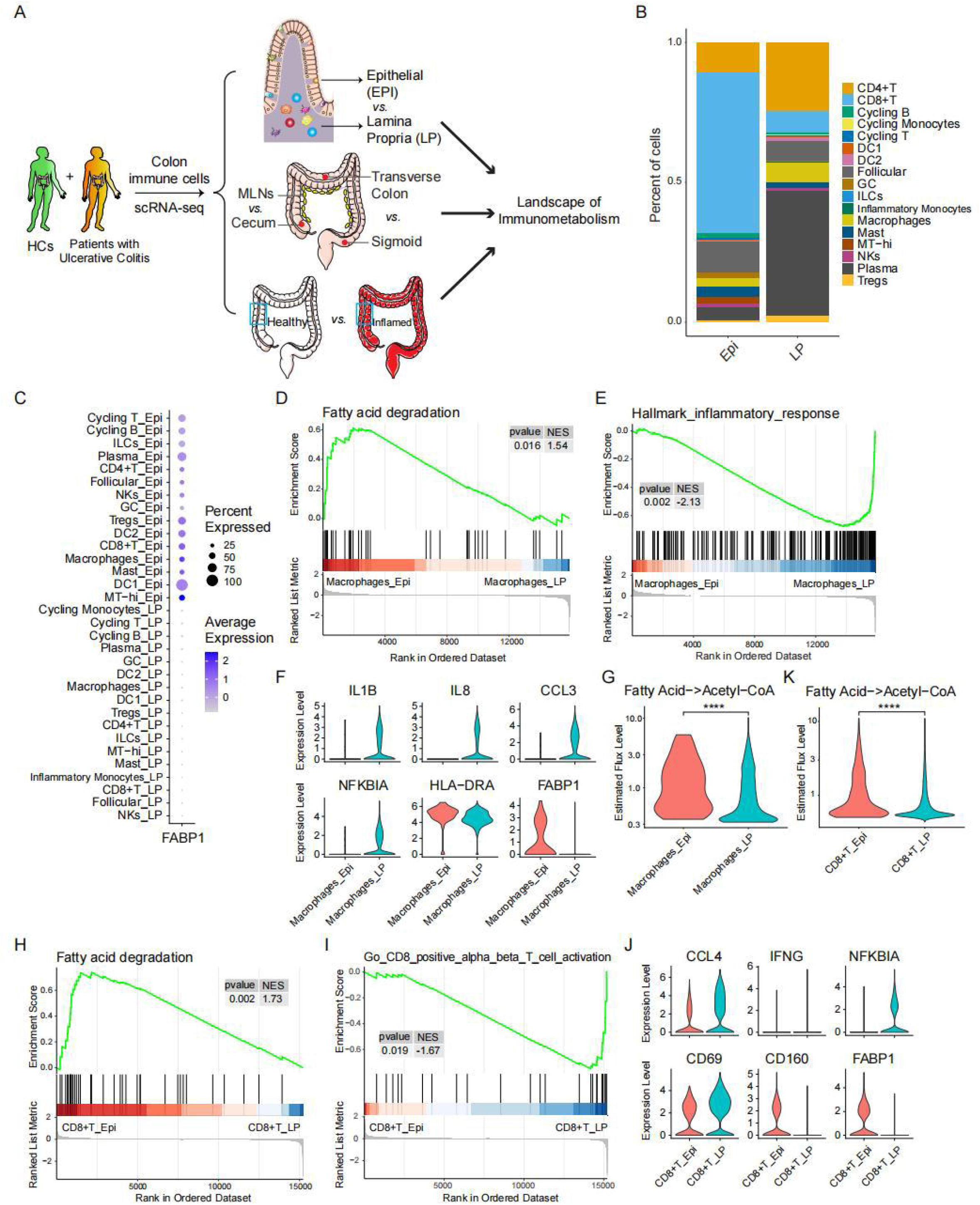
Distinct immune response in colonic epithelium compared with lamina propria. **(A)** A flowchart depicting the overall design of the study. **(B)** Bar plot shows the relative contributions of each immune cell subset in epithelium and lamina propria. **(C)** Dot plots show expression of FABP1 in each immune cell subset from epithelium and lamina propria. **(D-E)** GSEA analysis of indicated gene sets in macrophages from epithelium versus lamina propria of human colon. **(F)** Violin plots show expression levels of significantly differentially expressed genes in macrophages from epithelium versus lamina propria of human colon. **(G)** Violin plot shows relative level of indicated metabolic flux (estimated by scFEA) in macrophages from epithelium versus lamina propria of human colon. **(H-K)** GSEA analysis of indicated gene sets (H-I), violin plots show expression levels of significantly differentially expressed genes (J), and violin plot shows relative level of indicated metabolic flux in CD8^+^ T cells from epithelium versus lamina propria of human colon.

### Metabolically activated IgA+ plasma cells accumulated in lamina propria of sigmoid colon

After analyzing the Epi and LP of the human colon, we wished to study the differences among various parts of the colon. GSEA was performed to compare the 85 human metabolic pathways of each major immune subset (>50 cells in each group) from colon and MLNs. Interestingly, the results indicated that most of the metabolic pathways such as glycolysis, citrate cycle (TCA cycle), and oxidative phosphorylation pathways were upregulated in colonic IgA plasma cells compared with those in MLNs **(Figure S2A)**, which indicated colonic IgA plasma cells deserve to be explored in detail. According to existing findings, the IgA plasma cell is different from the cecum to sigmoid, which is significantly enriched in the sigmoid region, and a highly activated state of plasma cells in the distal colon compared with proximal colon plasma cells, characterized by greater accumulation, somatic hypermutation, clonal expansion (James et al., 2020a). In our study, we found that the metabolic score of B cell IgA plasma in sigmoid was significantly higher with the other two regions **(Figure 2A)**. And then we calculated the metabolic activity score of 85 pathways, which showed that the pathway activities interrelated with energy metabolism were higher in B cell IgA plasma from sigmoid, such as glycolysis, citrate cycle (TCA cycle), and pyruvate metabolism **(Figure 2B)**. The increased rate of energy metabolic pathways in sigmoid colonic B cell IgA plasma was further proved through the metabolic gene set signature scores **(Figure 2C)**, the fluxes results estimated by scFEA and scFBA **(Figure S2B, S2C)**, and GSEA **(Figure S2D)**. In addition to metabolic analysis, we also identified CD27 was highly expressed in sigmoid among other parts **(Figure 2D)**. As reported, IgA plasma cell could be divided into the CD27^+^ and CD27^-^ components naturally (Berkowska et al., 2011a). We explored that there was a significant up-regulation of overall metabolic score in the CD27^+^ group of IgA plasma **(Figure 2E)**, and many energy metabolic pathways were improved **(Figure 2F-G)**, which were consistent with metabolic fluxes predicted by scFEA **(Figure S2F)**. Therefore, the sigmoid colonic and CD27^+^ IgA plasma cells were metabolically activated compared with those in cecum and transverse colon and CD27^-^ group.

**Figure 2.**
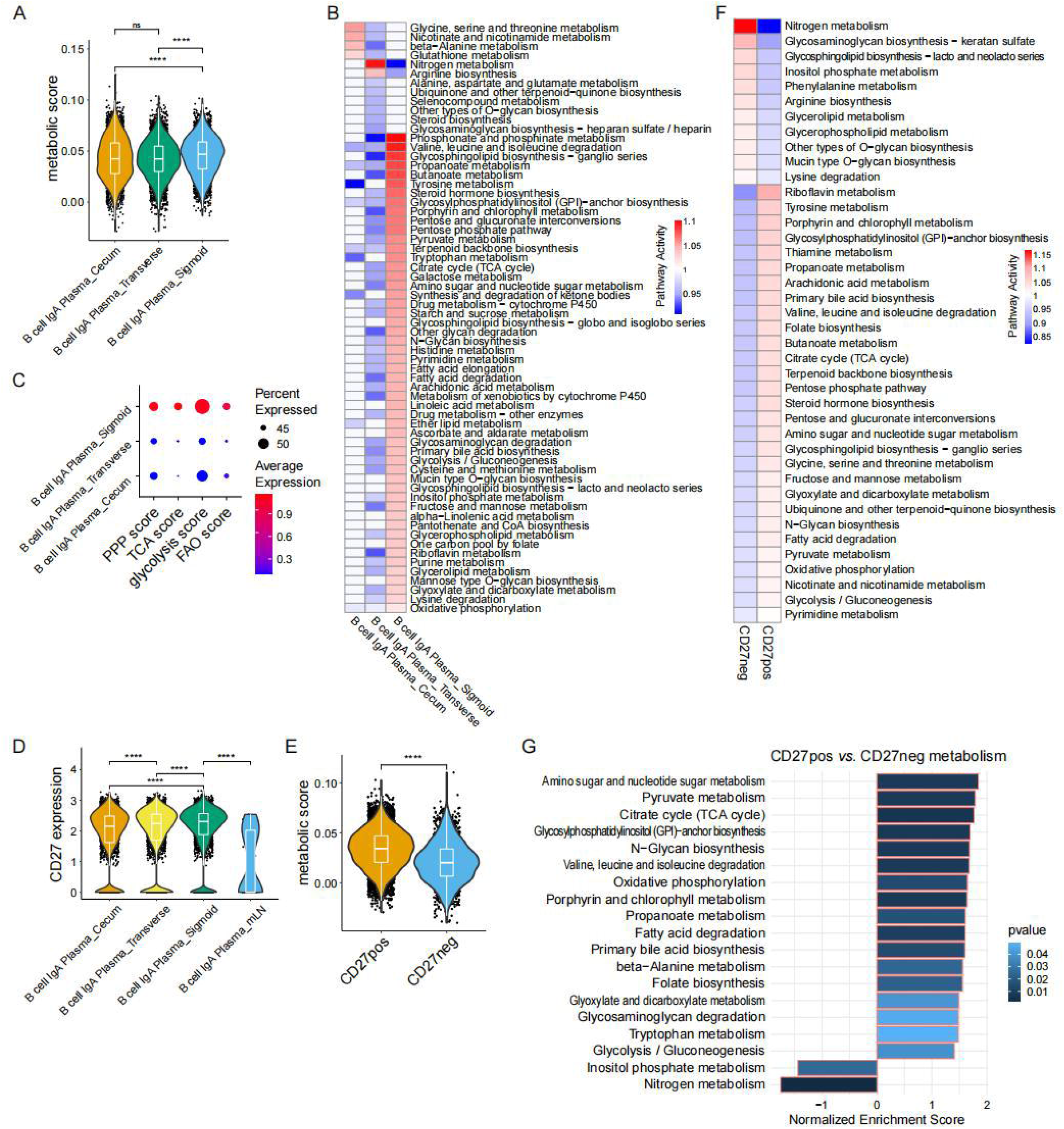
Metabolically activated IgA+ Plasma cells accumulated in lamina propria of sigmoid colon. **(A)** Violin plot shows relative level of metabolic score in IgA^+^ Plasma cells from cecum, transverse colon, and sigmoid of human colon. **(B)** Metabolic pathway activities in IgA^+^ Plasma cells from cecum, transverse colon, and sigmoid of human colon. Statistically non-significant values (random permutation test p>0.05) were shown as blank. **(C)** Dot plot shows relative level of indicated gene signature scores in IgA^+^ Plasma cells from cecum, transverse colon, and sigmoid of human colon. **(D)** Ditto plot shows expression of CD27 in IgA^+^ Plasma cells from cecum, transverse colon, sigmoid, and MLNs. **(E)** Violin plot shows relative level of metabolic score in CD27^+^ versus CD27^-^ IgA^+^ Plasma cells. **(F)** Metabolic pathway activities in CD27^+^ versus CD27neg IgA+ Plasma cells. Statistically non-significant values (random permutation test p>0.05) were shown as blank. **(G)** GSEA analysis of significantly enriched gene sets in CD27^+^ versus CD27^-^ IgA^+^ Plasma cells.

### Pro-inflammatory colonic macrophages were dysfunctional during ulcerative colitis

Besides the differences caused by the spatial factors, we wished to get knowledge of the alterations of immune cell during ulcerative colitis. We first identified that a lot of pathways related to inflammation that were upregulated in macrophages from the inflamed tissues of patients with ulcerative colitis versus those from healthy colons of HCs, such as TNF signaling, inflammatory response, and cytokine signaling **(Figure 3A)**, while some pathways which are of significance to macrophage functions, such as antigen processing and presentation, phagosome, and lysosome pathways were downregulated **(Figure 3B)**. Detailed analysis showed significantly elevated expressions of proinflammatory genes, such as S100A8, S100A9, S100A12 (positive regulators of macrophage inflammation (Xia et al., 2017)), FCN1 (a key driver of inflammatory macrophage state (Zhang et al., 2021)), CD44 (a cell-adhesion molecule playing crucial roles in inflammation (Puré and Cuff, 2001)), and HIF1A (a vital TF in glycolysis contributing to proinflammatory macrophages (Viola et al., 2019)). In contrast, the gene expression connected with metabolic process, and fatty acid oxidation, such as ATP6V1F, APOC1, APOE (key genes related to fatty acid metabolism and anti-inflammatory process in macrophages (Viola et al., 2019)), HLA-DPA1 (associated with antigen presentation ability (Ma et al., 2021)), LGALS3 (related to anti-inflammatory function (Dong et al., 2018)), and C1QC (known to enhance the M2 macrophages, anti-inflammatory macrophages (Benoit et al., 2012)) were downregulated **(Figure 3C)**. We reasoned that the macrophage might lose its original function and be more inclined to the pro-inflammatory macrophage. Our results also displayed an increased rate of glycolysis and pro-inflammatory response in macrophages from patients with ulcerative colitis **(Figure 3E-G, S3A)**, which showed no difference with previous findings that glycolysis had a positive effect on inflammatory macrophages (Viola et al., 2019).

**Figure 3.**
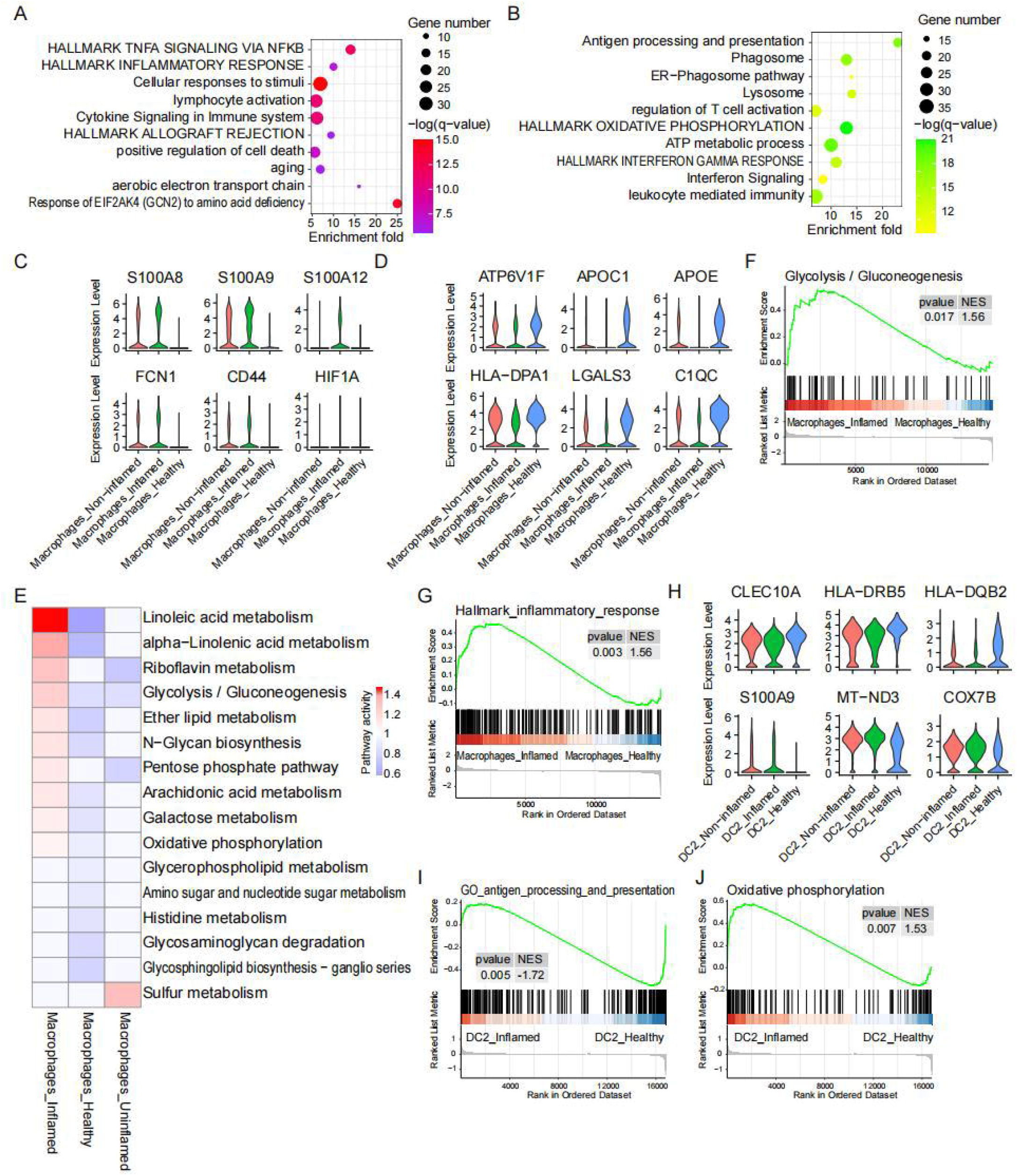
Myeloid cells from human colon were functionally dysregulated during ulcerative colitis. **(A-B)** Gene ontology analysis shows up-regulated pathways (A) and down-regulated pathways (B) in macrophages from inflamed tissues versus healthy ones. **(C-D)** Violin plots show expression levels of significantly up-regulated genes (C) and down-regulated genes (D) in macrophages from inflamed tissues versus healthy ones. **(E)** Metabolic pathway activities in macrophages from inflamed, non-inflamed, and healthy sections of human colon. Statistically non-significant values (random permutation test p>0.05) were shown as blank. **(F-G)** GSEA analysis of indicated gene sets in macrophages from inflamed tissues versus healthy ones. **(H)** Violin plots show expression levels of significantly differentially expressed genes in DC2 from inflamed tissues versus healthy ones. **(I-J)** GSEA analysis of indicated gene sets in DC2 from inflamed tissues versus healthy ones.

### The antigen presentation function of colonic dendritic cells was impaired in patients with ulcerative colitis

Another important myeloid immune subset with antigen processing and presentation ability was dendritic cell (DC). We identified that the DC2 from inflamed tissues reduced the expression of CLEC10A (a unique marker of CD1c^+^ DCs, (Heidkamp Gordon et al., 2016)), HLA-DRB5, and HLA-DQB2 that played essential roles in antigen processing and presentation of APCs **(Figure 3H)**. The GSEA also indicated that the antigen processing and presentation function was impaired in colonic DC2 from inflamed tissue compared with those from healthy ones **(Figure 3I)**. On the contrary, the expression of MT-ND3 and COX7B that correlated with the oxidation phosphorylation (OXPHOS) were higher in the DC2 from patients with ulcerative colitis than those from HCs **(Figure 3H)**, and the oxidative phosphorylation gene set was indeed more enriched in inflamed group **(Figure 3J, S3F)**. For the DC1, the results were similar with DC2 **(Figure S3B-E)**. We hypothesized that colonic dendritic cells became tolerogenic since it has been reported that OXPHOS was the main energy source of tolerogenic DCs, which possessed suppressed antigen presentation ability (Wei et al., 2020). Therefore, colonic myeloid cells, including the macrophages and dendritic cells, were dysregulated in patients with ulcerative.

### Lymphoid immune cells from human colon were functionally dysregulated during ulcerative colitis

In addition to colonic myeloid cells, we found that the regulatory T cells (Tregs) were activated during ulcerative colitis, upregulating TNFA signaling, OXPHOS, ATP metabolic process, and T cell activation pathways **(Figure 4A)**. The genes related to regulatory functions were elevated in Tregs from inflamed colon such as TNFRSF4 and TNFRSF18 (reduced the percentage of activated T cells and suppressed effector T cells (Zhou et al., 2020)), IL2RA (involved in Tregs development and function (Piotrowska et al., 2021)), CTLA4 (inhibited T cell activation by various suppressive functions (Mitsuiki et al., 2019)), BATF (regulated T helper cell differentiation (Park et al., 2018)), and MT-ND3 **(Figure 4B)**. We believed that improved regulatory function of Tregs confirmed by GSEA **(Figure 4C)** was correlated with increased rate of OXPHOS since mitochondrial energy metabolism strongly promote the function execution of Tregs (Chou et al., 2021). Furthermore, we identified the functionally impaired natural killer (NK) cells in patients with ulcerative colitis through **(Figure S4B)**. The genes had relations with NK functions were downregulated in NKs from inflamed tissue, such as XCL1 (encoded 2 chemokines known to recruit XCR1^+^ cross-presenting DCs into tumors (de Andrade et al., 2019)), GZMK (also known as Granzyme K, linked to NK function (Guo et al., 2010)), while the TGFB1 (an immunosuppressant for NKs (Mihi et al., 2014)) was upregulated **(Figure S4A)**.

**Figure 4.**
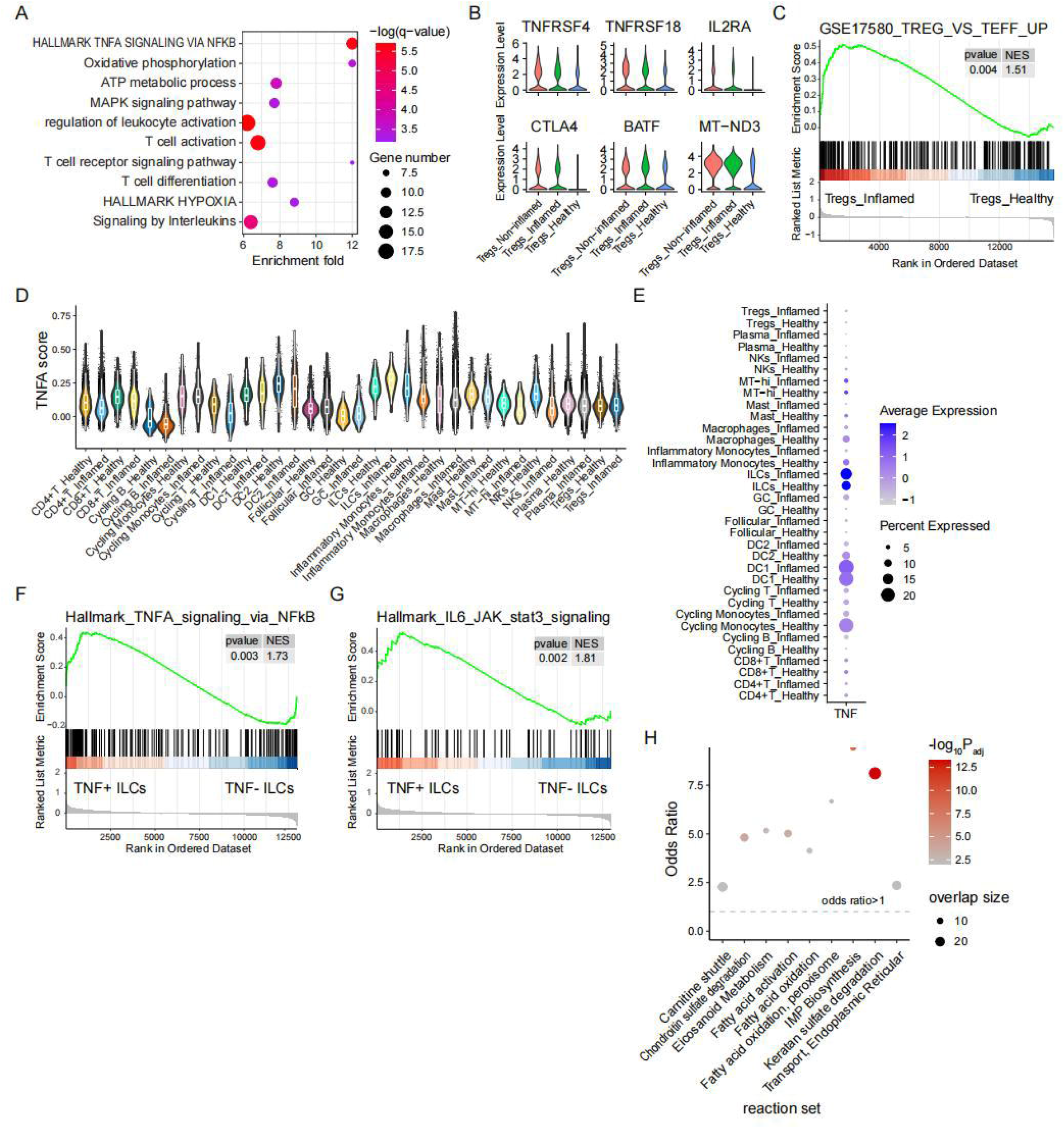
Colonic lymphoid immune cells became dysfunctional during ulcerative colitis. **(A)** Gene ontology analysis shows up-regulated pathways in Tregs from inflamed tissues versus healthy ones. **(B)** Violin plots show expression levels of significantly up-regulated genes in Tregs from inflamed tissues versus healthy ones. **(C)** GSEA analysis of indicated gene sets in Tregs from inflamed tissues versus healthy ones. **(D)** Ditto plot shows relative level of TNFA score in immune cells from inflamed tissues versus healthy ones. **(E)** Dot plots show expression of TNF in each immune cell subset from inflamed tissues versus healthy ones. **(F-G)** GSEA analysis of indicated gene sets in TNF^+^ versus TNF^-^ ILCs. **(H)** rMTA analysis was utilized to predict metabolic reactions whose knockout can further transform the metabolic state of the TNF^+^ ILCs back to the TNF^-^ ILCs. The significant metabolic pathways (FDR < 0.1) enriched by the top 10% MTA-predicted targets are shown. Y-axis represents the odds ratio of enrichment, and the horizontal dashed line corresponds to odds ratio of 1. The dot size corresponds to the number of enriched target reactions, and the dot color corresponds to -log_10_P_adj_. Half-dots plotted on the top border line correspond to infinity odds ratio values.

Anti-TNF therapy had the most significant effect for the treatment of ulcerative colitis (Pugliese et al., 2017), so we explored the TNF expression in each immune cell type. We found that not only did the innate lymphoid cell (ILC) have a high TNFA score but also it possessed higher expression of TNF at inflamed status **(Figure 4D-E)**. ILCs were then divided into the TNF^+^ ILCs and TNF^-^ ILCs, and TNF^+^ ILCs were indeed more pro-inflammatory **(Figure 4F-G)**. The robust metabolic transformation algorithm (rMTA, (Valcarcel et al., 2019)) was utilized to predict metabolic reactions whose knockout can reverse TNF^+^ ILCs to TNF^-^ ILCs metabolic changes to reduce its TNF expression. With the top 10% MTA-predicted targets enriched, potential target pathways, such as IMP Biosynthesis and Keratan sulfate degradation pathways, were identified that could be used to metabolically reprogram TNF^+^ ILCs **(Figure 4H)**.

### Prediction of metabolic targets for OSM+ inflammatory Monocytes

Because about 40% of patients do not respond to this treatment, only resting on the anti-TNF treatment to treat ulcerative colitis might be insufficient (Ben-Horin and Chowers, 2011). Then we defined a gene set called anti-TNF resistance genes using genes that come from the patients who do not respond to the treatment (Smillie et al., 2019c; Wang et al., 2016), and discovered that the inflammatory monocytes kept a high anti-TNF resistance score **(Figure 5A)**. Since the patients with nonresponse to the anti-TNF therapy significantly upregulated the OSM gene expression (Li et al., 2021), we subset the cluster of OSM^+^ inflammatory monocytes. The OSM^+^ inflammatory monocytes possessed a higher expression of G0S2 and PTGS2 (noted anti-TNF resistance genes (Kim et al., 2014; Smillie et al., 2019c)) and the genes associated with inflammation, such as IL1B, CXCL2, and NFKBIA **(Figure 5B)**. Consistently, the gene sets of inflammatory response and TNFA signaling were enriched in OSM^+^ inflammatory monocytes versus OSM^-^ ones **(Figure 5C-D)**. Using rMTA, the potential target pathways, including IMP Biosynthesis and OXPHOS, were enriched, which played crucial parts in reversing the OSM^+^ inflammatory monocytes to a normal state **(Figure 5E)**. As a consequence, we distinguished some potential metabolic reactions that can be combinate with anti-TNF therapy to treat ulcerative colitis.

**Figure 5.**
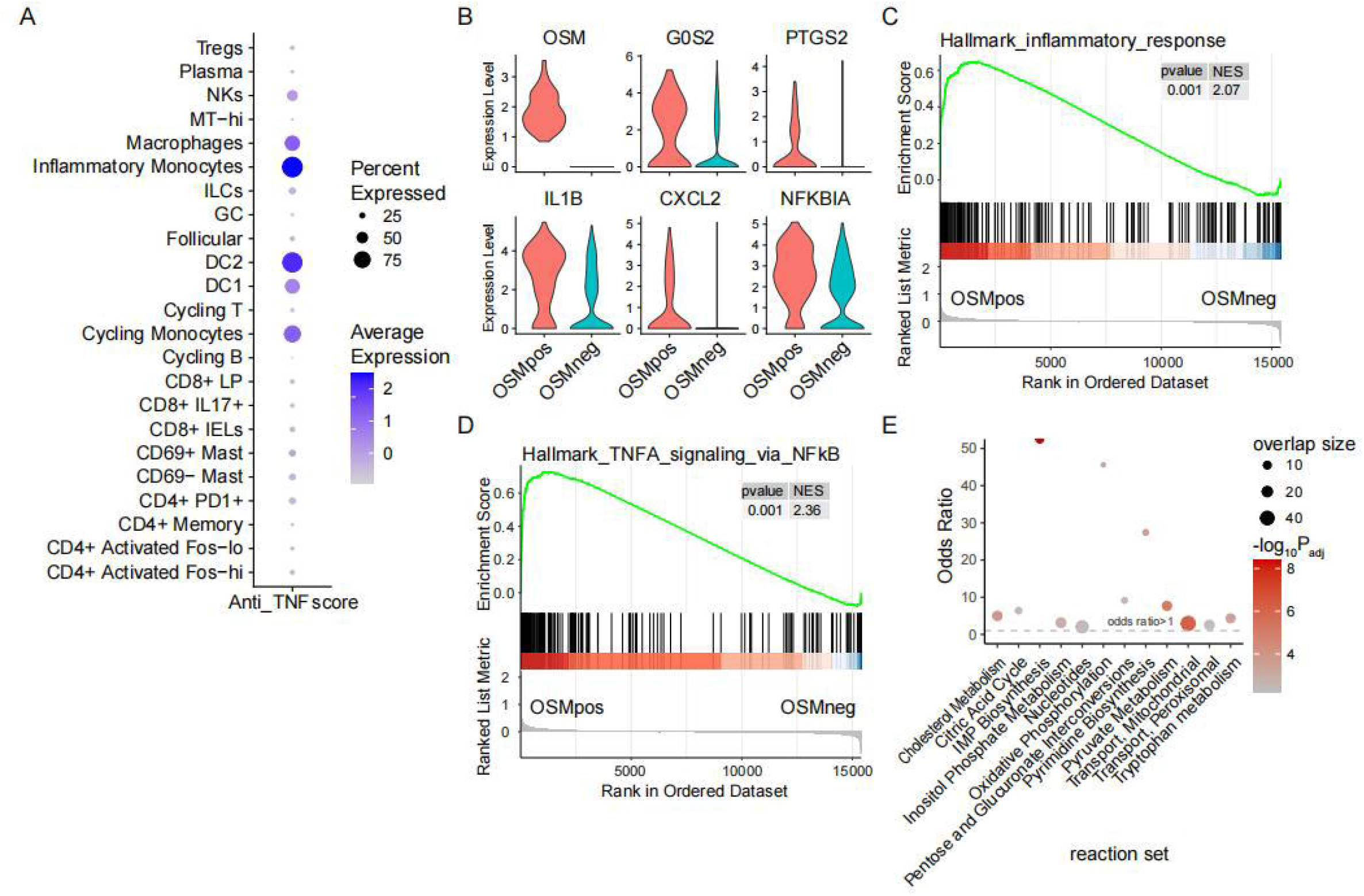
Prediction of metabolic targets for OSM^+^ inflammatory Monocytes. **(A)** Dot plot shows relative level of anti-TNF resistance score in each immune cell subset from inflamed sections of human colon. **(B)** Violin plots show expression levels of significantly up-regulated genes in OSM^+^ versus OSM^-^ inflammatory Monocytes. **(C-D)** GSEA analysis of indicated gene sets in OSM^+^ versus OSM^-^ inflammatory Monocytes. **(E)** rMTA analysis was utilized to predict metabolic reactions whose knockout can further transform the metabolic state of the OSM^+^ inflammatory Monocytes back to the OSM^+^ inflammatory Monocytes. The significant metabolic pathways (FDR < 0.1) enriched by the top 10% MTA-predicted targets are shown. Y-axis represents the odds ratio of enrichment, and the horizontal dashed line corresponds to odds ratio of 1. The dot size corresponds to the number of enriched target reactions, and the dot color corresponds to -log10Padj. Half-dots plotted on the top border line correspond to infinity odds ratio values.

## DISCUSSION

In this study, we depicted the immunometabolic mechanisms for the spatial immune specialization and the dysregulated immune response during ulcerative colitis at the single-cell level. These enabled us to map the metabolic alterations to distinct immune functions caused by spatial and disease-related factors. In doing so, we utilized the single-cell gene expression profiles of immune cells from different regions and states of the human colon (James et al., 2020b; Smillie et al., 2019a) to reveal their metabolic reprogramming by using some novel computational pipelines that were suitable for single-cell metabolism analysis.

We first found macrophages and CD8^+^ T cells in the Epi possessed a higher activity of fatty acid degradation, while they were less activated versus those in the LP **(Figure 1B-D)**. The up-regulation of FABP1 in the immune cells in the Epi might be the cause of an increased rate of fatty acid degradation since FABP1 has been reported to interact with PPARα, whose target genes include those encoding FAO enzymes (Gajda and Storch, 2015; Mallordy et al., 1995; Nakamura et al., 2014). Furthermore, it has been demonstrated that the M2 program (anti-inflammatory property) of macrophages requires the induction of fatty acid degradation (Huang et al., 2014; Odegaard and Chawla, 2013; Pearce and Pearce, 2013; Vats et al., 2006), and activated T cells could not engage in glycolysis but have an improved activity of fatty acid degradation on PD-1 ligation (Patsoukis et al., 2015). So, we hypothesized that the up-regulated fluxes of fatty acid degradation might be connected with the inactivated states of immune cells in Epi.

It was also very interesting to identify a cluster of metabolically activated IgA^+^ plasma cells in the sigmoid colon since there was no significant metabolic difference between IgA^+^ plasma cells in the cecum and transverse colon **(Figure 2A)**. We thought the increased rate of many energy metabolic pathways **(Figure 2B-C)** could favor the accumulation and functionality of IgA^+^ plasma cells in the human sigmoid colon. It has been reported that some metabolic pathways like glycolysis played very important roles in the proliferation and production of IgA antibodies of colonic IgA^+^ plasma cells (Caro-Maldonado et al., 2014; Dufort et al., 2014; Kunisawa, 2017). Furthermore, two human IgA^+^ B cell subsets have already been distinguished (Berkowska et al., 2011b): T cell-dependent CD27^+^IgA^+^ and T cell-independent CD27^-^IgA^+^ plasma cells. We identified that sigmoid colonic IgA^+^ plasma cells had higher CD27 gene expression **(Figure 2D)**, and CD27^+^IgA^+^ plasma cells indeed were more metabolically activated **(Figure 2E-G, S2F)**. The higher metabolic activity of sigmoid colonic IgA^+^ plasma cells might thus be related to CD27 expression.

Another important question we’d like to explore was whether the dysregulated metabolic reprogramming of human colonic immune cells led to the impaired immune response during ulcerative colitis, a subtype of inflammatory bowel disease (IBD) (Xavier and Podolsky, 2007). It’s well-known that the Hif1_α_-induced glycolysis and pentose phosphate pathway (PPP) were essential to the activation of inflammatory macrophages (Jha et al., 2015; Tannahill et al., 2013; Van den Bossche et al., 2017; Wang et al., 2017), so the improved glycolysis and PPP might contribute to the pro-inflammatory polarization of macrophages in the inflamed tissues of the colon **(Figure 3E-G, S3A)**. We also identified some genes of OXPHOS that were downregulated in the inflamed state **(Figure 3B-D)** (though some mitochondrion-coded genes were upregulated (data not shown)), and it has been demonstrated that OXPHOS was required for the induction of an M2 program (Van den Bossche et al., 2016; Vats et al., 2006). More importantly, we found downregulation of HLA-DR gene expression **(Figure 3H)** and suppressed antigen processing and presentation ability of human colonic dendritic cells (DC) in patients with ulcerative colitis versus HCs **(Figure 3I, S3C)**. Company with impaired immune function, the rate of OXPHOS was also increased **(Figure 3J, S3D-F)**, and this was a metabolic feature of tolerogenic dendritic cells (Sim et al., 2016).

For the lymphoid immune cells, the activated ATP metabolic process in colonic Tregs during ulcerative colitis might bring about improved regulatory functions **(Figure 4A-C)**, since key regulators of mitochondrial energy metabolism are required for optimal Tregs function (Beier et al., 2015). The activation of Tregs was considered to have a negative influence on the appropriate adaptive immune response. We also identified ILCs possessed a stronger TNFA signaling score and TNF gene expression among colonic immune subsets **(Figure 4D-E)**, and some potential target metabolic reactions were further found to reprogram TNF^+^ ILCs to a normal state **(Figure 4H)**.

Anti-TNF therapy is now well established as an effective therapeutic approach for ulcerative colitis. However, up to 40% of patients with ulcerative colitis or other IBD exhibit primary non-responsiveness to anti-TNF therapy (Ben-Horin and Chowers, 2014; Guerra and Bermejo, 2014). In our study, the anti-TNF resistance score was stronger in inflammatory monocytes compared with other immune cell types **(Figure 5A)**. The cluster of OSM^+^ inflammatory monocytes highly expressed other genes related to resistance to anti-TNF therapy, like G0S2 and PTGS2 (Kim et al., 2014; Smillie et al., 2019a) **(Figure 5B)**, and they were more pro-inflammatory than OSM-inflammatory monocytes **(Figure 5C)**. We also found disturbing the IMP, pyrimidine biosynthesis and OXPHOS might transfer the OSM^+^ inflammatory monocytes to OSM^-^ ones **(Figure 5E)**.

All in all, our investigation depicted a landscape of metabolic reprogramming of human colonic immune cells in different locations and disease states and further offered insights into treating ulcerative colitis by immunometabolic modulation.

## METHODS

### ScRNA-seq Data Availability and Data Processing

ScRNA-seq data of colon from healthy controls and patients with ulcerative colitis used in Figure 1 and Figure 3-5 can be accessed in Single Cell Portal (https://portals.broadinstitute.org/single_cell) under the accession number SCP259 (Smillie et al., 2019a). Data of human colon from different regions used in Figure 2 can be acquired in the Gut Cell Atlas (https://www.gutcellatlas.org/) (James et al., 2020b).

We excluded cells with gene number less than 200 or large than 6,000 (for data of human colon from different regions, is less than 700 or large than 6,000). Cells with mitochondrial gene percentage more than 20% were also removed in the study. The large filtered matrix was normalized using ‘LogNormalize’ methods in Seurat (version 3.2.2) (Stuart et al., 2019) with scale.factor = 10000. Variables ‘nCount_RNA’ and ‘percent.mt’ were regressed out in the scaling step and PCA was performed using the top 2,000 variable genes. Cell was annotated as reported previously (James et al., 2020b; Smillie et al., 2019a). Also, immune cell types including <50 cells were removed from the downstream analysis.

### Differential Gene Expression Analysis

The significant differentially expressed gene (DEG) lists were generated through Wilcoxon in Seurat (version 3.2.2) (FindMarkers function), and Logfc.threshold was set to be equal to 0.25, min.pct equal to 0.1, and adjusted p-Value less than 0.05. These DEGs were then utilized to perform enrichment analysis with Metascape platform (https://metascape.org) (Zhou et al., 2019).

Gene Set Enrichment Analysis (GSEA) (Subramanian et al., 2005) was also performed (nperm = 1000) with clusterProfiler R package (version 3.18.1) (Yu et al., 2012). The gene lists performed to GSEA were obtained through MAST in Seurat (version 3.2.2) (FindMarkers function); Lists of gene sets in KEGG database (http://www.kegg.jp) or MSigDB database (Bunis et al., 2020; Liberzon et al., 2015) (http://www.gsea-msigdb.org/gsea/msigdb/index.jsp) were used for further annotation.

### Gene Sets Score Calculation

To define the signature scores of specific signaling pathways, we used a list of metabolic genes from KEGG database, gene signatures termed “HALLMARK_INFLAMMATORY_RESPONSE”, “GO_CD8_POSITIVE_ALPHA_BETA_T_CELL_ACTIVATION”, “HALLMARK_TNFA_SIGNALING_VIA_NFKB” from MsigDB (Bunis et al., 2020), and gene set for resistance and susceptibility to anti-TNF blockade from a meta-analysis of 60 responders and 57 non-responders (Kim et al., 2014; Smillie et al., 2019a). These scores were then computed by using Seurat (AddModuleScore function) with default parameters.

### Prediction of potential target metabolic reactions with metabolic transformation algorithm

The significant DEG lists (obtained by MAST in Seurat) and the representative flux distribution (computed with iMAT (Shlomi et al., 2008)) of TNF+/- ILCs and OSM+/- inflammatory monocytes were used as inputs for the robust metabolic transformation algorithm MTA (rMTA) (Valcarcel et al., 2019; Yizhak et al., 2013) to predict target metabolic reactions whose knockout can transform cellular metabolic state from that of TNF+ ILCs and OSM+ inflammatory monocytes to that of the control states. The human Recon 1 GEM (Duarte et al, 2007) was selected for the MTA analysis, and the top 10% predictions were tested for significant overlap using Fisher’s exact tests. The detailed guides of using iMAT and rMTA could be found on GitHub (https://github.com/ruppinlab/covid_metabolism) (Valcárcel et al., 2019).

### Data Imputation and Normalization

To correct for the influence of the high frequency of dropout events caused by the low sequencing depth of scRNA-seq, scImpute algorithm (Li and Li, 2018) was utilized to impute genes with dropout rates large than 50%. In addition, to make it possible for comparison between different cell types and groups, we used deconvolution method (L. Lun et al., 2016). Briefly, we first utilized the scran R package (1.18.5) (Aaron et al., 2016) to calculate the size factor of each immune cell type annotated previously; And then, the normalized counts value of genes with dropout rate less than 0.75 were calculated by dividing counts by cell type-based size factor.

For immune cell types including >5000 cells, a training set containing 34 colon biopsies of 7 UC patients and10 HCs that has been described previously (Smillie et al., 2019a) was used to impute and perform the downstream analysis.

### Single-Cell Flux Estimation Analysis and Single-cell Flux Balance Analysis

To obtain single-cell flux panels of immune cells from different regions and disease states of human colon, two novel computational method, namely single-cell Flux Estimation Analysis (scFEA) Single-cell Flux Balance Analysis (scFBA) were utilized (Alghamdi et al., 2021b; Wagner et al., 2021). The imputed and normalized data of immune cells from colon were utilized as the input of scFEA and scFBA.

For scFEA, a graph neural network model was built to estimate cell-wise metabolic flux of colon immune cells based on the module gene file that contained metabolic genes for each module and the stoichiometry matrix that described relationship between compounds and modules, and they were available on GitHub (https://github.com/changwn/scFEA/tree/master/data). Also, the sc_imputation function was set to “False” since our data was imputed as mentioned above.

For scFBA, the Compass algorithm which can integrate our scRNA-seq profiles with prior knowledge of the metabolic network like Recon2 (Thiele et al., 2013) to infer metabolic states of immune cells (Wagner et al., 2021). We followed the instructions on how to install and use Compass that were available on GitHub (https://github.com/YosefLab/Compass). The developed tools were also utilized to postprocess and analyze the results of Compass (https://yoseflab.github.io/Compass/Compass-Postprocessing-Tutorial.html).

### Calculation of Metabolic Pathway Activity

A computational pipeline (Zhengtao et al., 2019) was further applied to the data with imputation and normalization to calculate the pathway activity score (https://github.com/LocasaleLab/Single-Cell-Metabolic-Landscape). Based on previous report, the average expression of all metabolic genes was first computed in each cell type; Then we transferred these absolute mean levels of gene expression to the relative expression across all cell types; The metabolic pathway activity score was further defined as the weighted average of all metabolic genes included in the specific pathway; Finally, we performed random permutation test to remove pathway activity scores that were not significantly higher or lower than average.

### Gene Expression Visualization

The bar plots, dot plots, violin plots, and heatmaps for percentage of cells and gene expression were generated using dittoSeq R package (version 1.2.4) (Bunis et al., 2020), Seurat (DotPlot and VlnPlot function), and pheatmap (version 1.0.12) (https://cran.r-project.org/web/packages/pheatmap). Data were auto-scaled with default parameters when using these packages.

### Statistics

In sections of differential gene expression analysis, GSEA, rMTA, scFBA, and of pathway activity calculation, p-value was computed with default methods of corresponding R packages. The Wilcoxon test was used to compare the difference of gene signature score, metabolic fluxes estimated by scFEA, and specific gene expression using ggpubr R package (version 0.4.0) (https://CRAN.R-project.org/package=ggpubr).

## ACKNOWLEDGMENTS

The numerical calculations in this paper have been done on the supercomputing system in the Supercomputing Center of Wuhan University. We thank the Supercomputing Center of Wuhan University for supporting on bioinformatic analysis.

## AUTHOR CONTRIBUTIONS

Q.Z. conceived and designed study, Q.Z. and H.Y. collected and analyzed data, Q.Z. and T.Z. wrote manuscript. All authors have approved the final version of this paper.

## DECLARATION OF INTERESTS

The authors declare no competing interests.

## FIGURE LEGENDS

**Figure S1. Distinct immune response in colonic epithelium compared with lamina propria, related to Figure 1**.

**(A-B)** Ditto plot shows relative levels of FAO and inflammatory scores in macrophages from epithelium versus lamina propria of human colon.

**(C-D)** Ditto plot shows relative levels of FAO and CD8^+^ T cell activation scores in CD8^+^ T cells from epithelium versus lamina propria of human colon.

**Figure S2. Metabolically activated IgA**^**+**^ **plasma cells accumulated in lamina propria of sigmoid colon, related to Figure 2**.

**(A)** Gene set enrichment analysis summarizing metabolic pathway alterations of indicated subpopulations in colon versus MLNs. A pathway was included in the heatmap if there was an alteration with p-value less than 0.05 in any cell type.

**(B)** Dot plot shows relative levels of indicated metabolic fluxes (estimated by scFEA) in IgA^+^ plasma cells from cecum, transverse colon, and sigmoid of human colon.

**(C)** Heatmap shows the compass-score differential activity of metabolic reactions (dots). Reactions are partitioned by Recon2 pathways (see Method) and colored by the sign of their Cohen’s d statistic. Nonsignificant correlations (BH-adjusted p < 0.1) shown in light color. Red dots indicated pathways are higher in sigmoid, while blue dots in cecum and transverse colon.

**(D)** GSEA analysis of indicated gene sets in IgA^+^ plasma cells from sigmoid versus cecum and transverse colon.

**(E)** Dot plot shows relative levels of indicated metabolic scores in CD27^+^ versus CD27^-^ IgA^+^ plasma cells.

**Figure S3. Myeloid cells from human colon were functionally dysregulated during ulcerative colitis, related to Figure 3**.

**(A)** Dot plot shows relative levels of indicated metabolic scores in macrophages from inflamed tissues versus healthy ones.

**(B)** Violin plots show expression levels of significantly differentially expressed genes in DC1 from inflamed tissues versus healthy ones.

**(C-D)** GSEA analysis of indicated gene sets in DC2 from inflamed tissues versus healthy ones.

**(E-F)** Metabolic pathway activities in DC1 and DC2 from inflamed and healthy sections of human colon. Statistically non-significant values (random permutation test p>0.05) were shown as blank.

**Figure S4. Colonic lymphoid immune cells became dysfunctional during ulcerative colitis, related to Figure 4**.

**(A)** Violin plots show expression levels of significantly differentially expressed genes in NKs from inflamed tissues versus healthy ones.

**(B)** GSEA analysis of indicated gene set in NKs from inflamed tissues versus healthy ones.

## Notes

The authors have declared that no conflict of interest exists.

### Competing Interest Statement

The authors have declared no competing interest.

