## Supplementary figures and images for "ScRNA-seq Identified the Metabolic Reprogramming of Human Colonic Immune Cells in Different Locations and Disease States"

### FigS1

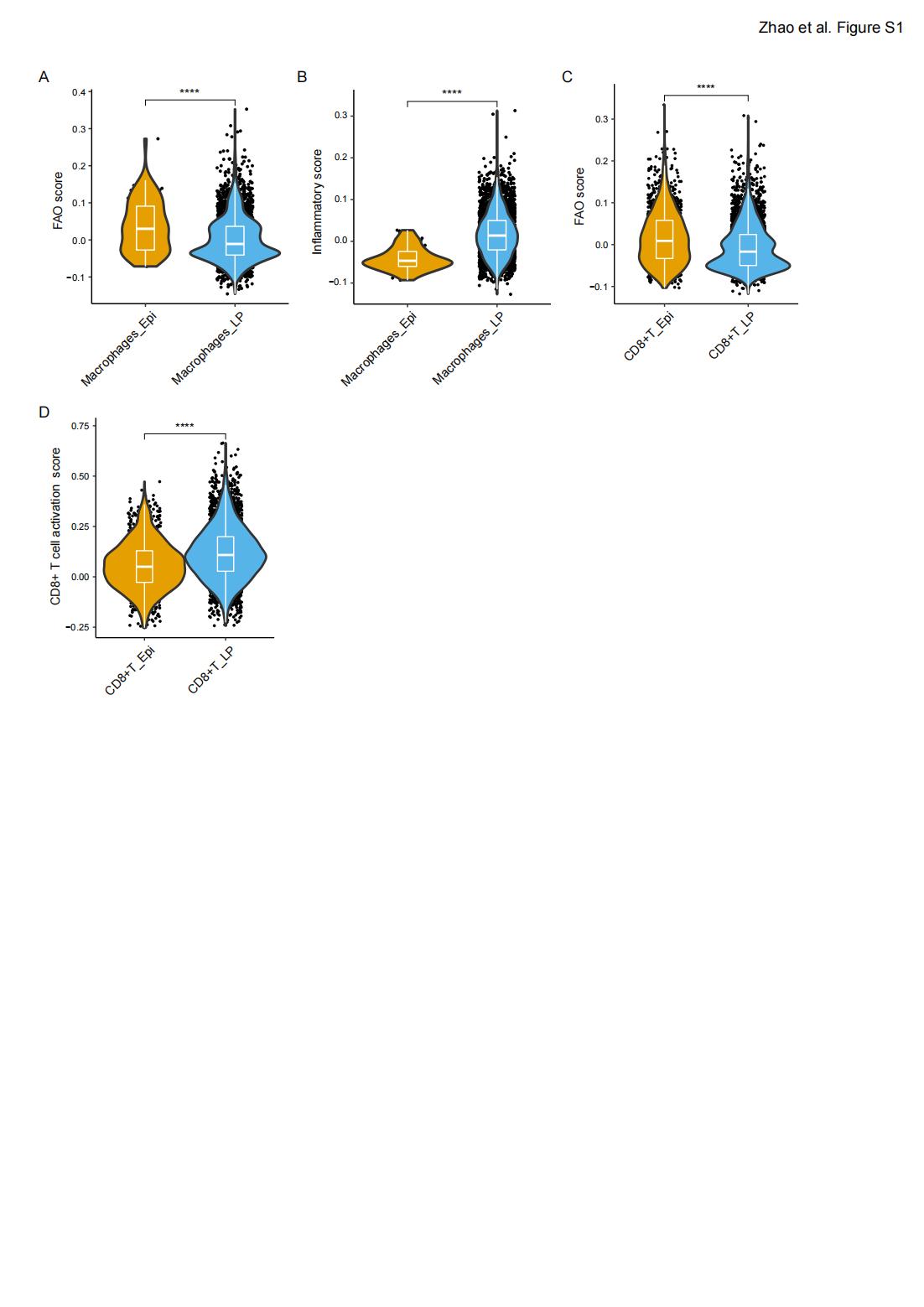

### FigS2

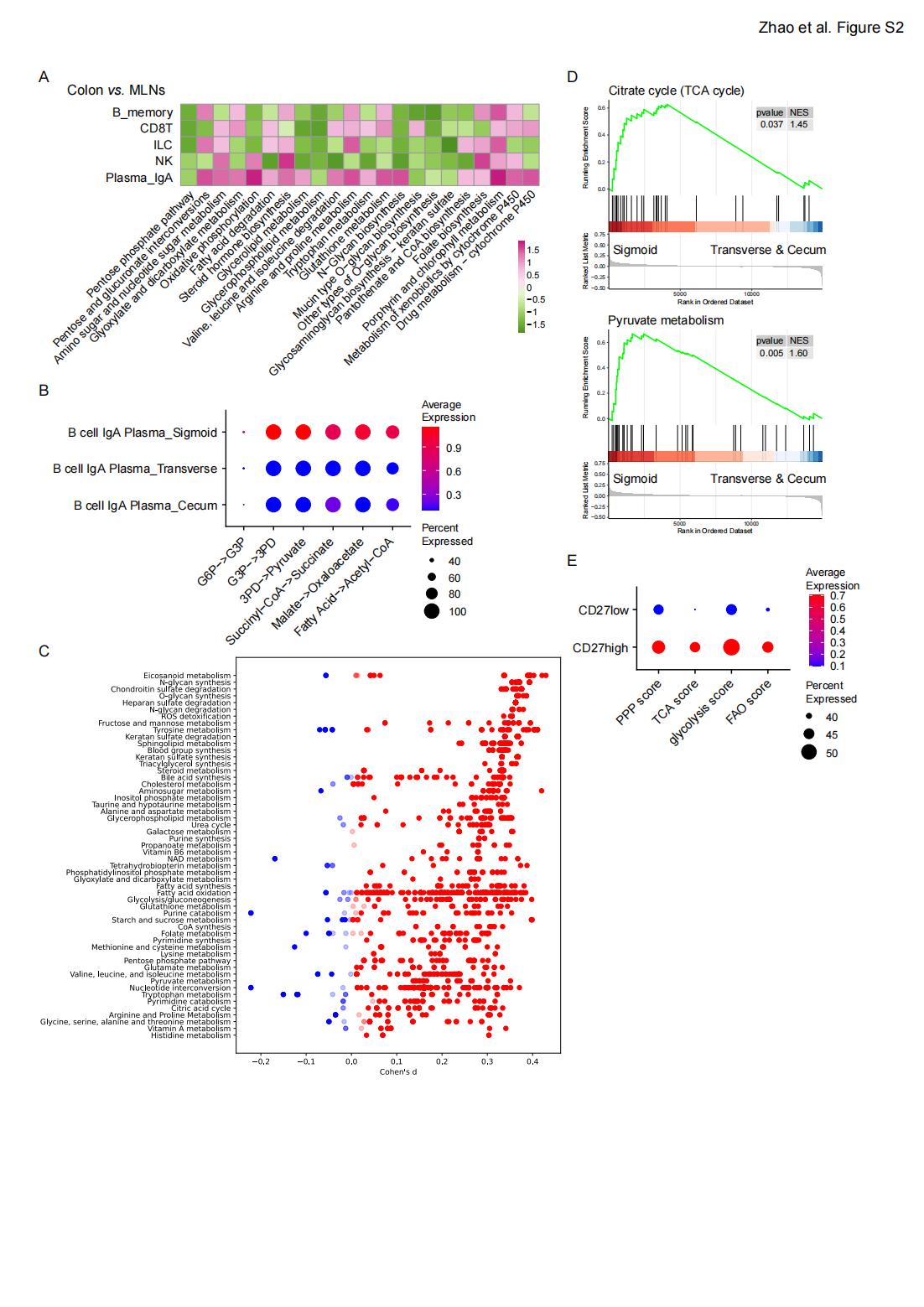

### FigS3

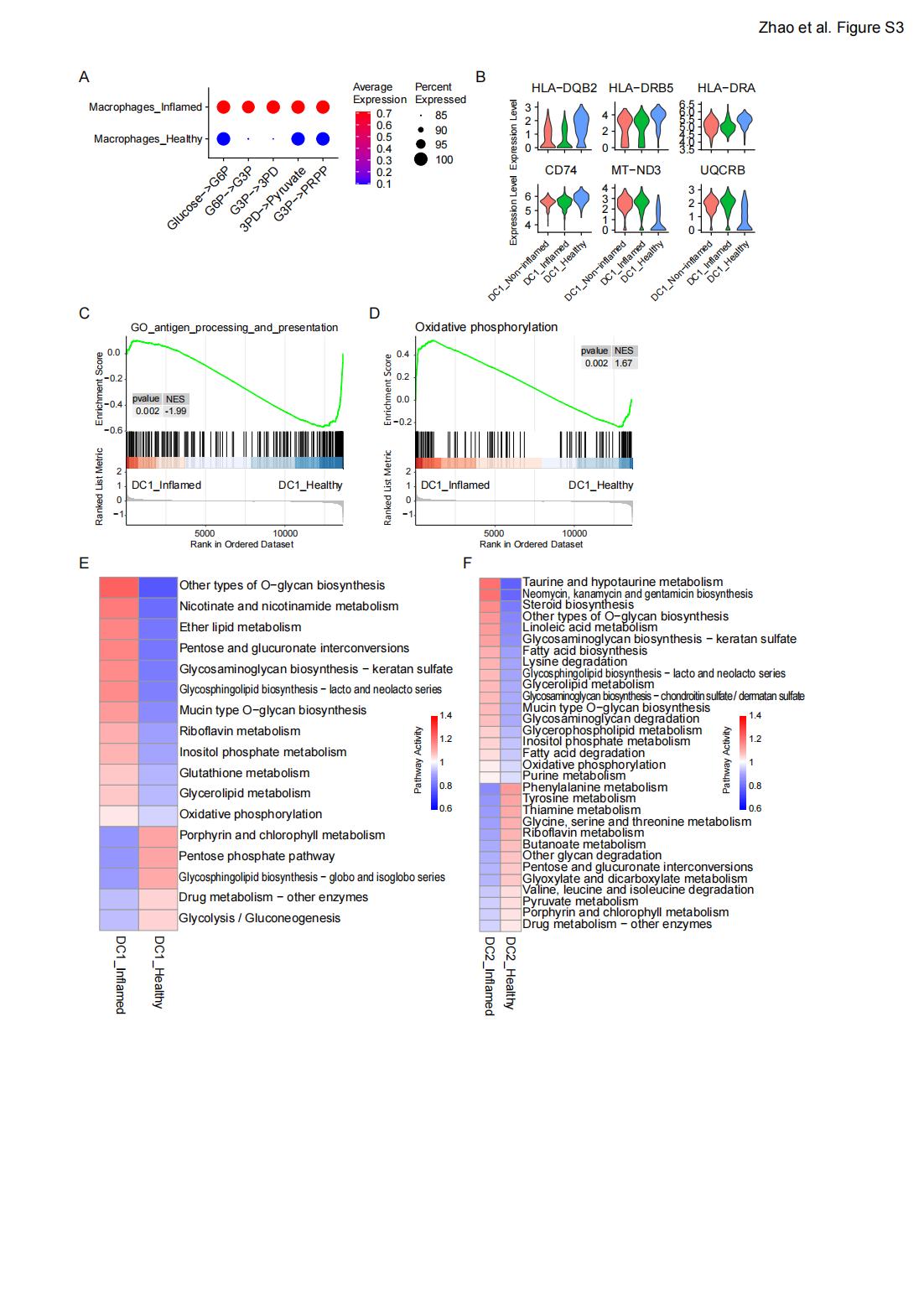

### FigS4

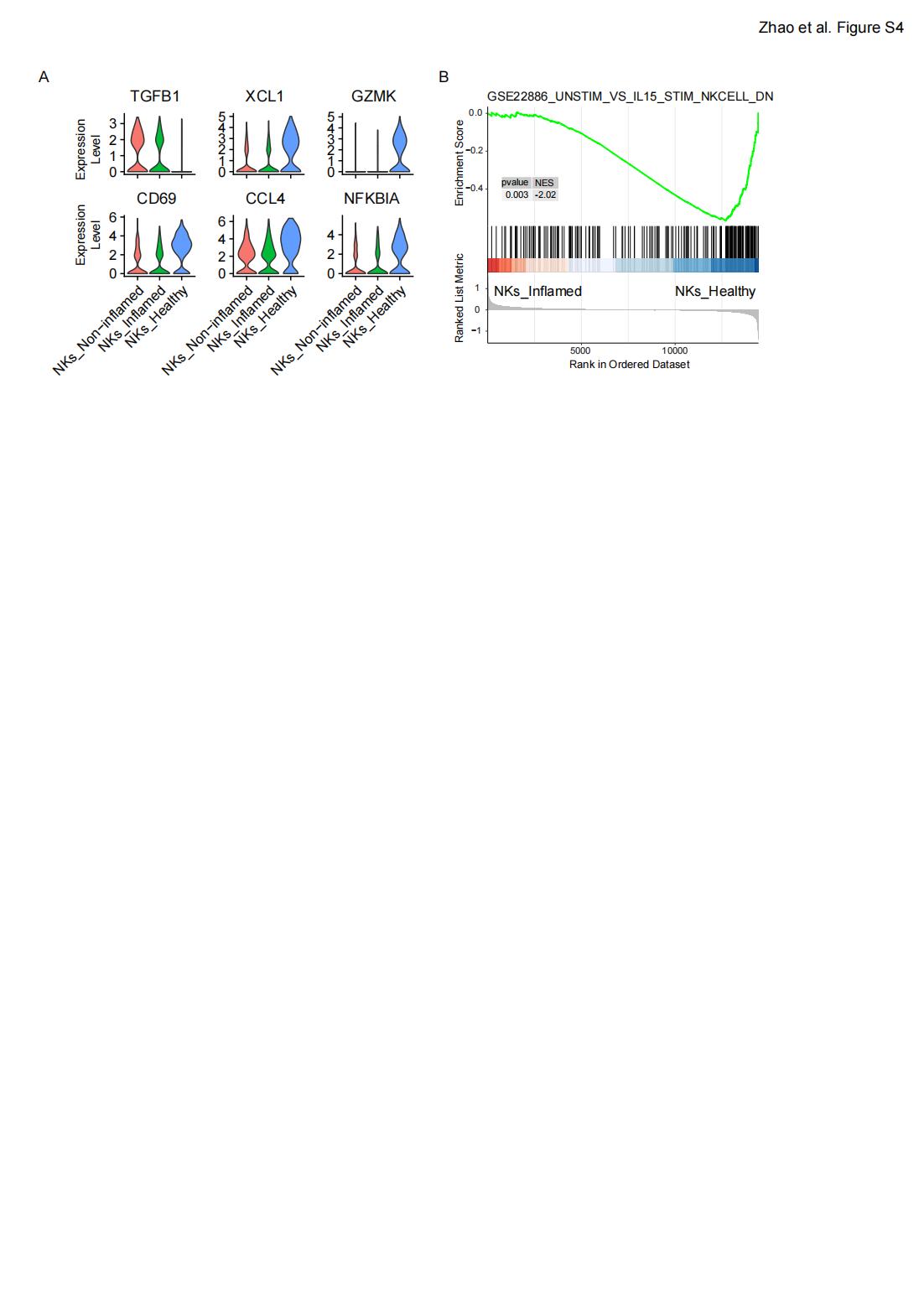
